# Sketching the SRA: experiment-wide similarity search across 4.8 million metagenomes

**DOI:** 10.64898/2026.10.07.757297

**Authors:** Roland Faure, Adrita Hossain Nakshi, Md. Hasin Abrar, Rayan Chikhi, Teo Lemane, Mohsen Taheri, Stephanie Won, Haonan Wu, David Koslicki, Paul Medvedev

**Affiliations:** Institut Pasteur, Université Paris Cité, CNRS UMR3525, Paris, France; Department of Computer Science and Engineering, Pennsylvania State University; LABGeM, Génomique Métabolique, CEA, Genoscope, Institut François Jacob, Université Paris-Saclay, CNRS, Evry, France; Institut Français de Bioinformatique, IFB-core, UAR 3601, CNRS, Villejuif, France; Department of Biochemistry and Molecular Biology, Pennsylvania State University; Huck Institutes of the Life Sciences, Pennsylvania State University; Department of Biology, Pennsylvania State University; One Health Microbiome Center, Pennsylvania State University

## Abstract

Recent efforts have demonstrated that systematic analysis of the millions of metagenomes in the Sequence Read Archive (SRA) enables groundbreaking biological discoveries. However, SRA-scale comparison of whole metagenomes has remained elusive. Here, we introduce the most comprehensive resource to date for experiment-to-experiment analysis of all SRA metagenomes, comprising FracMinHash sketches (470GB), hypergen sketches (19GB), and an all-pairs Jaccard similarity matrix (90GB), which we release online at https://github.com/RolandFaure/SRA-metagenome-sketches-databases. These resources enable a wide range of analyses, which we demonstrate with several applications. First, we use the FracMinHash sketches to uncover temporal trends, finding that the amount of newly discovered microbial sequence continues to grow, rather than saturate or shrink. Second, we use the hypergen sketches, which enable nearest-neighbor search, to identify co-assembly candidates and thereby improve assembly quality. Third, we use the similarity matrix to analyze all SRA metagenomes, identifying duplicated submissions and over 50K candidates for metadata correction. Together, our resources unlock systematic analysis of metagenomic data at unprecedented scale.

## 1 Introduction

The Sequence Read Archive (SRA) is the central public archive for sequencing data, containing millions of experiments and spanning a wide range of organisms, environments, experimental designs, and sequencing technologies [19]. Because the SRA stores primary sequencing data, it captures sequence diversity that is not necessarily represented in assembled-genome resources such as GenBank [30]. Metagenomes in particular are rarely exhaustively analyzed due to their complexity. Recent work shows that conventional genome-centric resources are missing substantial microbial diversity that is nevertheless present in large collections of public metagenome sequencing data [26]. A metagenomic experiment samples the genetic material of a whole microbial community (or a targeted subset of gene(s) in the case of amplicon experiments) and the raw sequencing data can uncover information about community composition, functional potential, environmental context, and strain-or population-level genetic diversity [7]. Therefore, the SRA remains an unparalleled primary resource for metagenomic analysis.

Several recent large-scale efforts have demonstrated that systematic analysis of SRA experiments can yield important biological discoveries (i.e. Serratus [8], Logan [7], MetaGraph [17], and Pebblescout [33]). These tools provide *sequence-to-experiment* comparison, whereby the user provides a short nucleotide sequence as a query and is returned all matching SRA experiments. While these are powerful tools, they do not allow efficient *experiment-to-experiment* comparison, i.e. comparing whole experiments against each other. As we demonstrate in this paper, such comparisons can facilitate a wide range of applications such as co-assembly, duplication detection, and metadata correction.

The size and structure of the SRA make experiment-to-experiment analysis difficult. Metagenomic experiments in the SRA (identified by unique *accessions*, used here synonymously with sequencing datasets) often correspond to large collections of short, fragmented reads, and the full collection of all 4, 774, 811 metagenomic accessions is too large (1.2 PB) to support naïve all-vs-all analysis. Previous work has shown that the sketching technique could make analyses at this scale tractable [5, 28]. The Logan project [7] made substantial headway by extracting and compressing all the non-singleton 31-mers within each accession. Nevertheless, even this compressed representation requires 184 TB, making all-vs-all analysis of SRA metagenomes cost-prohibitive.

In this work, we compute and compress a sketch database of all SRA metagenomic data (Figure 1). The sketch database is composed of two layers: FracMinHashes [16], taking 470 GB, and hypervectors, generated by the hypergen sketching technique [25, 38], taking 19 GB. Using the hypervector database, we then compute an all-pairs Jaccard similarity matrix. We provide two versions of the matrix: a full one storing all similarities above 5% (1.1 TB) and a filtered one storing only the neighbors with similarity either above 20% or in the top 10,000 of the closest similarities (90 GB). The sketch database and matrix are publicly hosted and freely available for download through our GitHub landing page. We provide several interfaces to the data, including command-line tools and a web-based interface that enables users to find all accessions similar to their own experiment.

**Figure 1:**
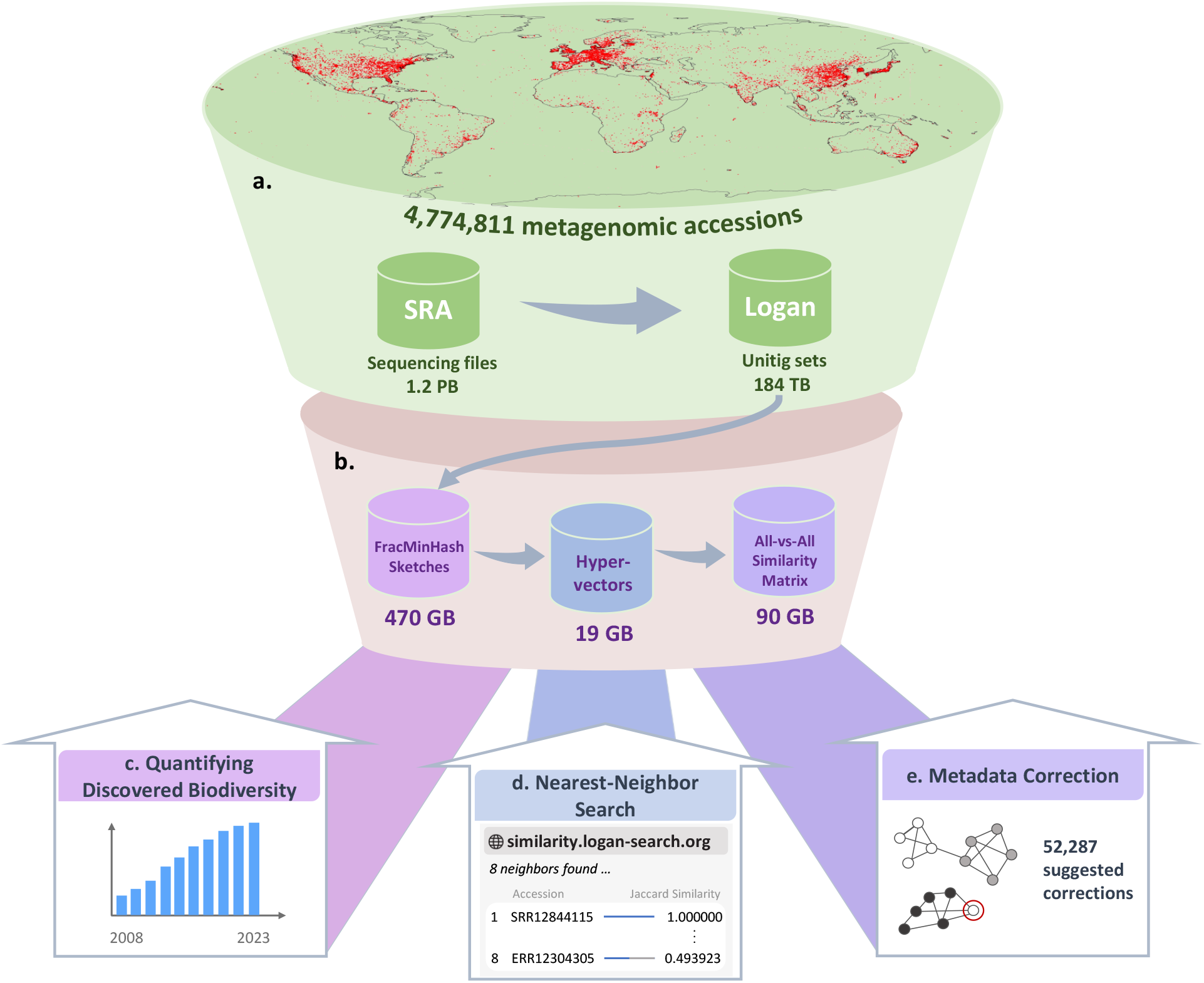
New databases for the large-scale analysis of SRA metagenomes. **(a)** Map showing the distribution of the metagenomes and the size of the corresponding sequencing and Logan unitig databases. **(b)** Description of the three resources we construct from these metagenomes. Left: the FracMinHash database contains a FracMinHash sketch of each accession”s non-singleton 31-mers. Center: the hypervector database contains a hypervector representation of each accession”s FracMinHash sketch. Right: the similarity matrix stores Jaccard similarity estimates between all pairs of accessions. **(c)** Application of the FracMinHash database: quantifying the metagenomic diversity over time. **(d)** Application of the hypervector database: a web service that allows to find similar SRA accessions to any user-provided dataset. **(e)** Application of the all-vs-all similarity matrix: correcting the SRA metadata by detecting metadata outliers. Not shown is an application to detect duplicate submissions.

Our sketch database and similarity matrix provide the most comprehensive-to-date resource for experiment-to-experiment analysis of all SRA metagenomic data. To demonstrate its potential impact, we explore several possible applications. First, we show how the FracMinHash database can be used to characterize *k*-mer diversity and novelty over time, finding that the amount of newly discovered microbial sequence continues to grow, rather than saturate or shrink. Second, we show how the hypervector database can help identify candidates for co-assembly when provenance fails to do so, thereby improving assembly quality. Third, we show how the similarity matrix can help uncover clusters of duplicated datasets in the SRA, helping flag potentially problematic submissions for further investigation. Finally, we show how the matrix can be used to identify and correct missing or erroneous SRA metadata, paving the way for automating more stringent metadata checks during the submission process.

This work also introduces a key methodological advance. Sketching naturally leads to downstream errors in the Jaccard similarity estimates. However, the error of the hypergen sketching approach (introduced in [38]) was only tested on a small set of genomes which do not represent the full diversity of SRA metagenomic datasets. Moreover, a brute-force simulation approach covering the vast space of possible datasets is computationally infeasible. To overcome this, we develop an algorithm that provably reduces the simulation problem to sampling from binomial distributions. This provides a rigorous characterization of the error associated with our Jaccard estimates and enables us to calculate bias (which turns out to be negligible) and derive confidence intervals for all possible inputs. Moreover, our algorithm allows us to tune the dimension of our hypervectors so as to achieve the desired balance between estimator accuracy and database size. This methodological innovation enables us to deploy the estimator at the scale of the SRA while maintaining accuracy guarantees.

## 2 Results

The Sequence Read Archive (SRA) contains 4,774,811 metagenomic accessions and 1.2 PB of raw metagenomic sequencing data (using the Logan v1.1 freeze, corresponding to December 2023). The Logan project reduced each accession to the set of its non-singleton 31-mers, stored compactly as unitigs and taking a total of 184 TB [7]. Though the unitig representation is already a substantial reduction in size, it is still prohibitive for most experiment-to-experiment analyses. In order to broaden the types of questions that can be asked of the data, we compute two new types of reduced representations of each accession: a FracMinHash and a hypergen sketch (Figure 1). Using the hypergen sketches, we also compute an all-pairs similarity matrix for all 4.8M accession pairs. Further summary statistics are in Sup. Table 1. All data can be accessed through our resource landing page^1^ and is stored for free and easy download access via PSU ScholarSphere and AWS. In the following, we describe each component of the sketch database and matrix together with its applications.

### 2.1 FracMinHash sketch database

FracMinHash (FMH) is a widely used technique [27] to sketch genomic data in a way that retains the ability to do a wide-range of downstream analyses. We construct a database of FracMinHash sketches of every accession (using a sampling rate of 1/1000) and develop a custom compression scheme to store it (Section 4.7 and Sup. Note 7). Our scheme reduces the size of the database down to 470 GB, a 6-fold reduction compared to 3.1 TB in its native Sourmash [16] format and a 390-fold reduction compared to 184 TB in Logan unitigs.

Figure 2(a) shows the size of the FMH in every accession, i.e. the number of its canonical 31-mer hashes^2^. The distribution breaks down by assay type, with 79% of accessions characterized as AMPLICON and 13% characterized as WGS (i.e. whole genome sequencing), with the remaining accessions being various other assay types (such as RNA-Seq, WGA, etc.). AMPLICON accessions have substantially smaller sketches (median 346) than WGS (median 70,039), consistent with the targeted, low-complexity nature of amplicon protocols such as 16S rRNA gene or ITS profiling, where the number of distinct genomic sequences (and hence distinct *k*-mers) is inherently limited by the amplified locus. WGS accessions yield markedly more variable sketches (i.e. WGS has interquartile range 8,455–254,761 compared to 148–715 for AMPLICON), reflecting higher sequencing depth across diverse genomic backgrounds. The heterogeneous “Other” category has a median (344) that is nearly identical to AMPLICON, indicating that the majority of non-WGS, non-AMPLICON accessions are also derived from targeted or low-complexity sequencing experiments such as clone-end sequencing assay types. However, “Other” has a much longer upper tail, likely driven by high-coverage protocols like Hi-C. Overall, these observations indicate that because FracMinHash sketches mirror the diversity of the underlying experiments, any SRA-wide metagenomic analysis is best stratified by assay type and, even within an assay type, must account for drastic variability of sketch sizes.

**Figure 2:**
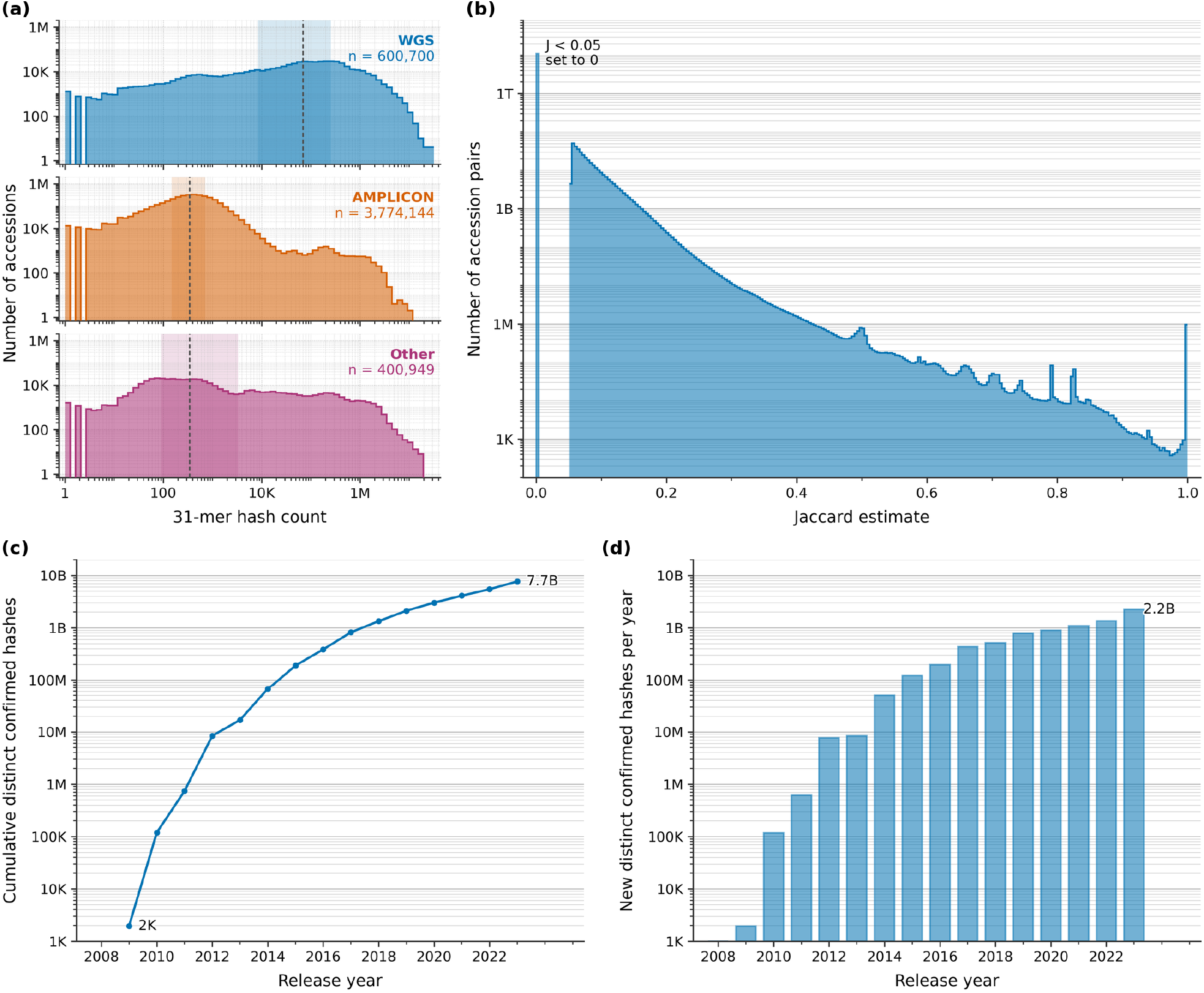
Scale, similarity structure, and temporal accumulation of diversity for 31-mer FracMinHash sketches across all SRA metagenomic accessions. **(a)** Distribution of 31-mer FracMinHash sketch sizes, stratified by sequencing assay type. Each subpanel shows the histogram of sketch sizes, i.e. the number of distinct canonical 31-mer hashes per accession, partitioned into whole-genome shotgun sequencing (WGS), amplicon-based sequencing (AMPLICON), and all remaining assay types combined (Other). Bins are uniformly spaced on a log_10_ scale spanning the full observed range (1–32,106,901 hashes per accession), and counts are shown on a log_10_ *y*-axis. Dashed vertical lines indicate the per-category median and shaded bands span the interquartile range. **(b)** Distribution of Jaccard estimates in our full all-pairs similarity matrix of all 4.8M SRA metagenomes (log_10_ *y*-axis). Bins are the 256 levels of the 8-bit quantization used to store the matrix; pairs estimated below 0.05 were set to zero at construction and form the single bar at *J* = 0, leaving (0, 0.05) empty. Self-similarity values of 1 are excluded. The small peaks between 0.5 and 0.9 likely reflect the quantized Jaccard values inherent to comparing datasets with few hashes. **(c)** Cumulative number of distinct canonical 31-mer confirmed hashes observed through the end of each calendar year (log_10_ *y*-axis); the total grew from 1,946 in 2009 to approximately 7.7 billion by the end of 2023, a roughly six-order-of-magnitude expansion over fifteen years. **(d)** Number of new distinct hashes confirmed in each calendar year (hashes are credited to the second year they were observed; log_10_ *y*-axis); annual novelty increased nearly monotonically from ~ 2K in 2009 to ~ 2.2B in 2023, with no sign of saturation. Callouts in c and d annotate the endpoints of each series (c, 2008 and 2023; d, 2023) in thousands (K), millions (M) or billions (B) as appropriate.

#### 2.1.1 Characterizing *k*-mer diversity and novelty over time

Given the global accumulation of sequencing data, one might ask if we have begun to exhaust the space of all existing microbial sequence. The metagenome *k*-mers in the SRA can help answer this question, as they represent an ongoing, distributed sampling of microbial sequence space. In particular, what fraction of the 10^18^ possible canonical 31-mers have been discovered in microbial data and has that number begun to saturate over time? Our FMH sketch database allows us to answer this question quickly, because the number of distinct canonical 31-mers is approximately 1000×the number of distinct hashes.

An immediate concern is that sequencing errors may generate 31-mers that appear twice in an accession but do not correspond to microbial sequence. To account for this, we say that a canonical 31-mer is “confirmed” if it has appeared in at least two calendar years, and we say that its “year of confirmation” is the second year in which it appears. Errors due to technical artifacts such as errors during PCR amplification are largely filtered out because they are unlikely to appear in two separate experiments. Sequencing errors are even less likely to be confirmed, as this would require that the same spurious sequence must arise in at least two independent reads within one accession, and then in at least two independent reads again in an accession from a different year. This four-concordant-error event has a probability that scales with the fourth power of the sequencing error rate and hence the effect of *k*-mers attributable to sequencing error becomes negligible. In summary, by only tracking 31-mers that are confirmed, we screen out most of the possible sequencing errors or technical artifacts that occur in a single sample.

As expected, the cumulative total of confirmed hashes continues to grow (Figure 2(c)), with 7.7B confirmed hashes as of 2023, corresponding to approximately 7.7T canonical 31-mers. More interestingly, the number of new hashes confirmed each year (Figure 2(d)) also continues to grow (as opposed to saturating or decreasing), representing continued increasing discovery of sequence diversity not previously represented in the SRA. We also see that certain environments, like soil, continue to yield a high rate of new hashes per megabasepair of sequencing while other environments, like the human gut, experience falling rates of novelty over time (Sup. Fig. 1; Sup. Note 6.1). This confirms what has previously been observed, that we have begun to exhaust the microbial sequence content of the human gut, but are nowhere close to exhausting the microbial sequence content of soil [26, 24, 1, 21].

Overall, these observations are consistent with two complementary interpretations: microbial sequence space remains far from exhausted even at present sequencing scales, and/or ongoing microbial evolution continuously generates novel sequence variants that are captured each year as new metagenomes are sequenced and deposited. While it is outside the scope of our analysis to weigh these interpretations against each other, the FMH sketch database allows this type of analysis to be done in a way that previous representations of SRA metagenomes (i.e. 184 TB of unitigs) do not.

### 2.2 Hypervector database

For applications where *k*-mer-specific information is not needed, FracMinHash sketches are unnecessarily large and cumbersome. We therefore transform each FMH sketch into a much smaller 4,096 byte hypervector using DotHash [25] projection (Section 4.1, Figure 3a). This combination of FracMinHash and DotHash procedures is known as hypergen sketching [38]. Our hypervector database takes 19 GB in total, a 24-fold reduction of the FMH database.

**Figure 3:**
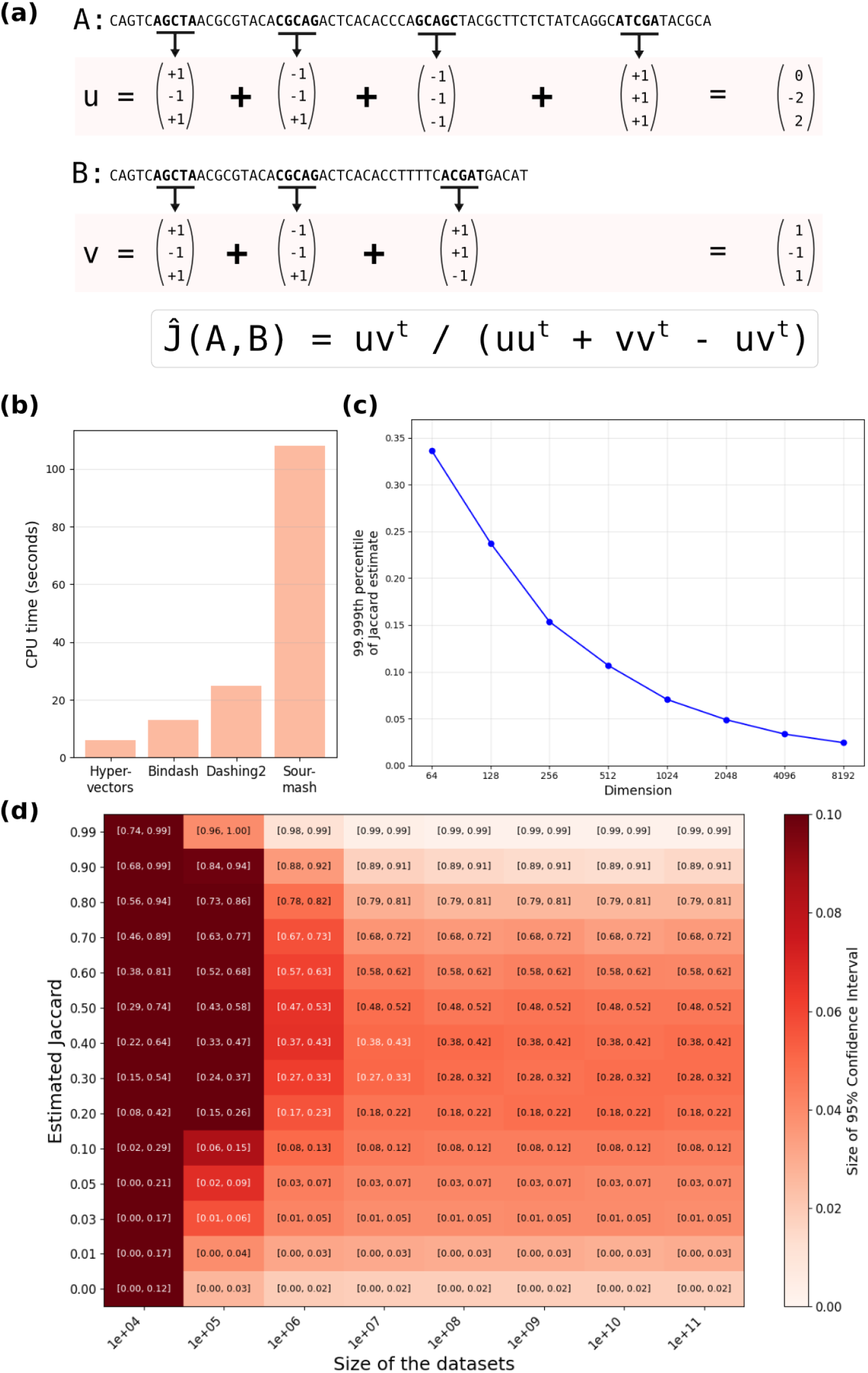
Hypervector-based Jaccard estimator. (a) Toy example illustrating how hypervectors are computed and used to estimate Jaccard similarities. In this example, *k* = 5 and *d* = 3. (b) All-versus-all Jaccard distance estimation times for four different software, including ours (“hypervectors”), on a subset of 3,537 datasets. Sketching times are not included. For BinDash [39] and Dashing2 [3], the sketches were parametrized to be the same size (4,096 bytes) as the hypervectors. Details provided in Sup. Note 6.3. (c) Upper 99.999 percentile of the Jaccard estimation distribution for pairs of 1-million *k*-mers datasets with true Jaccard similarity 0, plotted for different choice of hypervector dimension. (d) 95% confidence intervals of the Jaccard estimator, with each cell showing the interval and colored according to its width. We focus on pairs of datasets of equal sizes, which is the most difficult case for the estimator. The intervals were computed using our novel algorithm in Section 4.3.

We use the hypervectors to quickly estimate the Jaccard similarity of a dataset against all other accessions in a database (Figure 3b, Section 4.1). Note that other tools exist for estimating Jaccard similarity, but our preliminary results indicated that the hypervector approach exhibited the most promise for scaling up to the required sizes (Sup. Fig. 2c, Sup. Note 6.3). We therefore implemented a search engine where researchers can quickly compare their dataset against our hypervector database. After generating a hypervector of their dataset (which can be either an assembly or a collection of reads), users can use our free web service (https://similarity.logan-search.org) to quickly find all neighbors with similarity above a user-specified threshold; alternatively, they can request a user-specified number of nearest neighbors. Both the generating and querying processes typically complete within 30 seconds.

The estimated Jaccard values differ from the true ones due to the randomness introduced by the hypergen sketching processes. We evaluate the nature and scale of this error using a novel and provably correct sampling algorithm, using it to construct confidence intervals around our estimates (Section 4.2). First, we find that the error is provably independent of sequence content, conditioned on the true Jaccard similarity and the number of *k*-mers in each accession. We therefore compute error as a function of these three parameters; in particular, we compute error statistics for dataset sizes in the range of 10^4^ to 10^10^ *k*-mers and Jaccard similarities in the range of 0 to 0.99. We choose 10^4^ as the lower cutoff as it captures 98% of all SRA metagenomes. Second, we measure the bias of our estimator. We find that the absolute bias is always *<* 0.001 on the tested ranges, implying that it is negligible, especially since the true Jaccard is itself an imprecise measure of true biological similarity. Third, we observe that, on the tested ranges, the root mean square error is highest when the datasets are of equal sizes (Sup. Note 6.2), consistent with DotHash theory [25]. Fourth, focusing on tested datasets of equal sizes, we observe that the confidence intervals are generally widest for smaller sizes and Jaccard values closer to 0.5 (Figure 3d). For datasets with at least 1M *k*-mers (corresponding to approximately 26% of all accessions and 97% of all WGS accessions), the width of the 95% confidence interval is in the worst case 6 percentage points. As expected, this error prohibits the use of our hypervectors for fine-scale resolution (e.g. separating similarities of 2% and 4%); however, many analyses do not need such a level of resolution. Fifth, the accuracy of the estimator becomes unreliable for datasets containing fewer than 10,000 *k*-mers (corresponding to approximately 2% of all accessions). Sixth, we break down the source of the error and find that, consistent with theoretical expectations, the FMH sketch is generally the dominant source for smaller datasets while DotHash is generally the dominant source for larger datasets (Figure 2). Seventh, we calculate the Jaccard estimate cutoff at which the non-zero similarity detection false positive rate for datasets of at least 1 million *k*-mers drops below 10^−5^. In other words, for datasets with true Jaccard of zero, the estimated Jaccard will be above this threshold no more than 1 in 100,000 cases. By plotting it as a function of the hypervector dimension (Figure 3c), we identified *d* = 2048 as the smallest dimension at which this threshold becomes 0.05 and made that the dimension of our hypervectors.

#### 2.2.1 Identifying co-assembly helper samples using the hypervector database

Co-assembly is the process of improving the assembly of a sample by combining its reads with those of a biologically similar helper sample. Though it can risk contaminating the assembly, the risk is often acceptable given that co-assembly can enhance genome completeness by increasing effective coverage of shared taxa, particularly for low-abundance organisms [22, 10, 29]. Traditionally, co-assembly helper candidates have been chosen from the same study and/or the same biome; however, such samples are not always good choices as their sequence content can vary greatly, especially for complex communities such as soil microbiomes. Instead, researchers can use our nearest-neighbor search to find a helper sample based on sequence similarity from across the entire SRA metagenomic corpus.

To understand the potential for using our nearest-neighbor search for improving co-assemblies, we select 14 accessions as targets for improvements. For each target, we refer to its most similar WGS SRA-wide accession as the *SRA-wide helper* and its most similar accession in the same BioProject as the *project helper*. Six of the accessions are chosen arbitrarily from among BioProjects with only one public sequencing run (i.e. there was no project helper), and eight are chosen from accessions that had project helpers but the similarity to the project helper was less than to the SRA-wide helper. We assemble each target individually, together with its SRA-wide helper, and together with its project helper (when available). We leave out 10% of reads for later validation and downsample each helper so that its read volume is capped at that of the target. We then compare the assemblies based on the number of bases in long (≥10kbp) contigs that are supported by at least one of the held-out reads mapping with a read mapping quality ≥10. Further details are in Section 4.5.

Figure 4 shows that SRA-wide helpers can in most cases greatly improve assembly quality, in terms of useful bases assembled. When project helpers are not available, the SRA-wide helper improves the target-only assembly in five out of six cases, with a median improvement of 40% more bases. For the 14 targets overall, using a helper improves the assembly in 13 of the cases. Among these, the SRA-wide help was a better choice that the project helper in 11 cases, with a median improvement of 20% in these cases. These results demonstrate the potential of using our nearest neighbor similarity search to help improve helper-identification for co-assembly.

**Figure 4:**
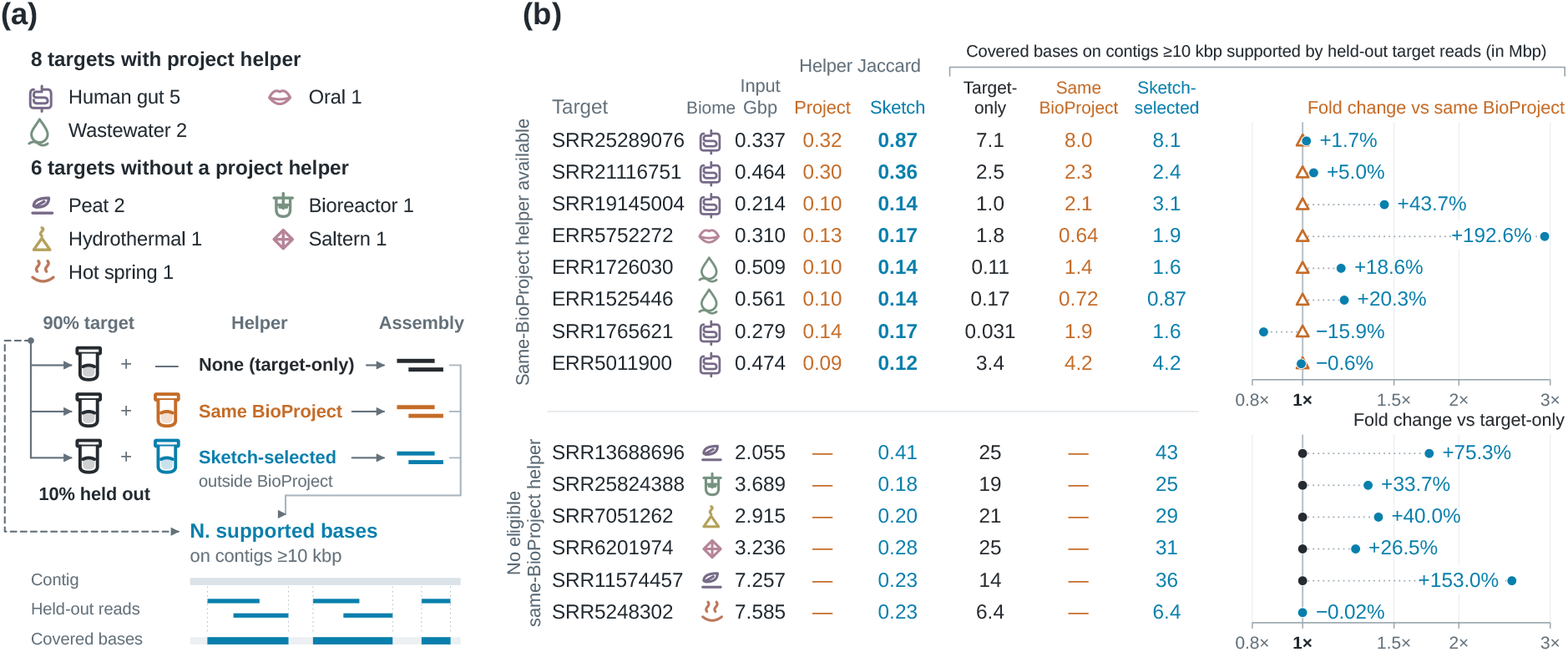
Co-assembly with hypervector-database-selected helpers improves assembly quality. “Input (Gbp)” is the number of bases used for the target-only assembly. “Helper Jaccard” refers to our estimated Jaccard similarity of the target to the helper. We count the number of bases on contigs ≥ 10 kbp that are covered by at least one alignment of a held-out target read with mapping quality ≥ 10. The last three columns of the table shows this count in assemblies based on just the target (“Target-only” column, and black dots), the target mixed with the most similar helper from the same BioProject (“Same BioProject” column, and orange triangles), and the target mixed with the most similar SRA-wide helper (“Sketch-selected” column, and blue dots). The right-most fold-change plot visualizes these counts relative to each other.

### 2.3 All-pairs similarity matrix

Using our hypervector database, we compute a matrix of Jaccard similarity estimates (Section 4.1) between all 4.8M metagenomes. To achieve a computation of this scale, we rely on the powerful Eigen linear algebra library [13] and store the results in a custom compression format (Section 4.7). Following the error experiments previously described (Figure 3(c)), Jaccard estimates below 0.05 are removed in order to control the non-zero similarity detection false positive rate. This results in a matrix of size 1.12 TB. Further analysis showed that the mean number of neighbors per accession is 145,982 and that the vast majority of pairs had similarity less than 0.2 (Figure 2(b)). To make a more nimble version of the matrix, we considered additional filters to remove uninformative information (Sup. Table 4). We ultimately decided on a filter that retains all neighbors with similarity *>* 0.2 and the top 10,000 neighbors, while discarding everything else. This trims excessive neighborhood lists while guaranteeing that no neighbors that are similar are removed. The resulting filtered matrix is an order-of-magnitude smaller at 90 GB; the mean number of neighbors is 8,486.

Figure 2(b) shows that the number of entries in our (full) matrix that have a specific Jaccard value generally increases as the values decreases, with the major exception of a high number of entries with a Jaccard of one. In general, it is not surprising that a large fraction of accessions have at least one neighbor with non-zero estimated Jaccard. A significant portion of microbial genomes contain broadly conserved regions (e.g. 16S, 18S, and ITS regions), resulting in hashes that are highly shared, e.g. there is a FMH hash that appears in 48% of all FMH sketches, and another nine hashes that each appear in 29%-46% of all sketches.

#### 2.3.1 Identifying duplicated and potentially problematic datasets

The distribution of Jaccard similarity estimates in the all-pairs matrix showed a pronounced peak at one (Figure 2(b)). This motivated a more detailed analysis of hv-duplicate accessions, which we define as an accession that has at least one other accession with an estimated similarity of 1 and hence the same hypervector. An obvious reason for this might be that the accessions have very small amounts of data. We therefore identified 151,405 hv-duplicate accessions whose raw sequencing data was larger than 10 Mbp, corresponding to 3% of all accessions. Most hv-duplicate accessions occur as simple pairs, i.e. they are associated with only one other matching accession. However, we also observed a small number of larger hv-duplicate groups i.e. a group of accessions with the same hypergen sketch.

We investigated the reasons behind these hv-duplicate groups. First, some are formed from accessions with very small FMH sketches (e.g. five hashes), suggesting that the high similarity is driven by low-quality or uninformative reads. For example, this accounted for the largest hv-duplicate group, containing 333 accessions and spanning multiple BioProjects and `organism` labels. Second, some of these groups are due to duplicated uploads within the same project. For example, this accounted for the largest hv-duplicate group among WGS samples, which contained 106 accessions from the same BioProject (`PRJEB60243`). The raw read files for these accessions were completely identical, suggesting that the same data had been uploaded multiple times, either by mistake or by acceptable reuse across studies. Third, some of these groups spanned multiple BioProjects and seemingly unrelated sample types. For example, accessions from a Congo cave study (`PRJNA982691`, `SRR24958364`) and a maize study (`PRJNA821718`, `SRR18673646`) appeared in the same hv-duplicate group, and manual inspection confirmed that their sequence files were identical. This may represent repeated and potentially problematic submissions. Overall, our all-pairs similarity matrix allows us to identify many cases of duplicated SRA accessions; in the future, our hypervector database could be used to automate such checks as part of the SRA submission process.

#### 2.3.2 Identifying and correcting metadata issues

Metadata is manually entered during submission into the SRA and is consequently susceptible to human error [18]. Submitters frequently enter incorrect information or use overly general terms. However, reliable metadata is of particular importance, especially as it will become central to training AI methods in the near-future.

We first relied on exploratory analysis to evaluate the potential of using our similarity matrix to systematically correct metadata. For instance, accessions ERR11146382 and ERR11146401 carry only the generic “metagenome” `organism` tag, yet all of their top 100 nearest neighbors have similarity of at least 0.8 and are labeled “mouse gut metagenome,” strongly suggesting that they should be relabeled.

Another example is BioProject PRJEB45799, whose 2,394 accessions are labeled as “human gut metagenome.” Principal component analysis (PCA) of its hypervectors, however, show what appear to be two distinct clusters (Figure 5A). The meaning of these clusters becomes evident when we projected the hypervectors from another BioProject (PRJNA662107) onto the same plot. PRJNA662107 contains well-annotated human oral and gut metagenomes that mapped distinctly to the two observed clusters. It is likely that during upload of PRJEB45799, all accessions were mistakenly labeled with the same “human gut metagenome” tag, even as some of them corresponded to an oral metagenome, a hypothesis confirmed by the corresponding article [35].

**Figure 5:**
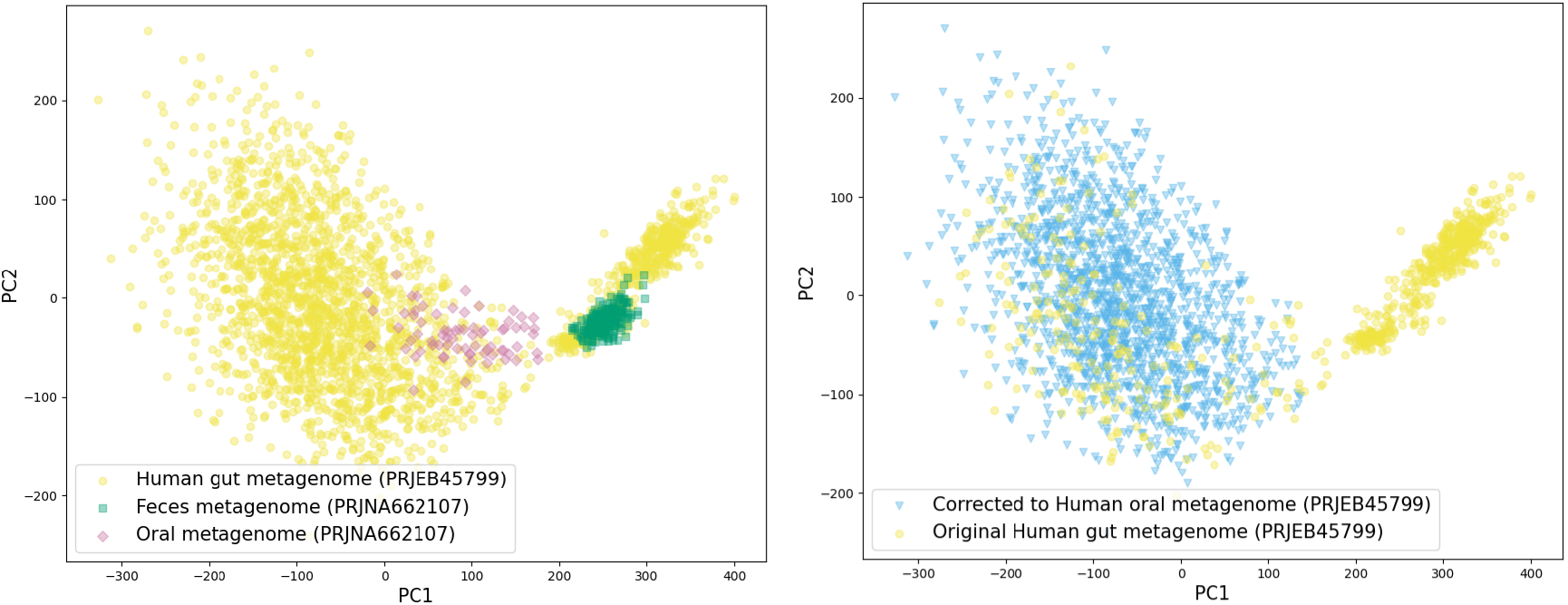
Principle component analysis of hypervectors of project PRJEB45799, showing likely metadata mislabeling. The two axes represent the two principle component axes computed from the hypervectors corresponding to accessions in project `PRJEB45799`. The left panel projects the hypervectors of `PRJEB45799` (yellow, all initially tagged as “human gut metagenome”) as well as the hypervectors from project `PRJNA662107` on these axes (with green corresponding to “feces” tag and pink corresponding to “oral” tag). The right panel shows the `PRJEB45799` hypervectors but colors them according to our proposed correction. The analysis supports that while all accessions of `PRJEB45799` are labeled as “gut” in the SRA, many of them actually correspond to “oral”.

Our exploratory analysis indicates that the all-pairs similarity matrix can provide a quantitative mechanism to systematically identify and relabel metadata fields by leveraging the consensus labels of an accession”s nearest neighbors. Using this idea, we develop a proof-of-concept strategy to relabel the `organism` metadata tag in all 600,700 WGS accessions (Section 4.6 describes the details of this strategy). Our strategy results in the proposed relabeling of 52,287 (8.7%) `organism` metadata entries (listed in a Supplementary File). The majority of modifications (34,866 cases) involve refining non-specific “general metagenome” labels to the more precise “human gut metagenome” label. There were also 1,680 corrections where human gut metagenomes were reclassified as oral metagenomes, with most of these (1,628) coming from the aforementioned BioProject PRJEB45799.

We relied on spot-checking to validate the accuracy of the proposed corrections. For BioProject PRJEB45799, all the corrections were consistent with the PCA analysis (Figure 5B), though some accessions that lie in the “oral metagenome” cluster were not relabeled. To spot-check other proposed relabelings, we randomly selected 30 accessions from distinct BioProjects, comprising 20 cases of refined tags (i.e. relabeling to a more specific tag) and 10 cases of corrected tags (i.e. relabeling to a tag that is inconsistent with the existing tag). Manual verification against the corresponding publications revealed that 24 of the 30 relabelings (80%) were accurate (Sup. Table 3). The remaining six cases had relabeling suggestions which reflected the fact that highly similar microbial communities could be associated with different biomes. For example, one correction relabeled a “mouse gut metagenome” as “human gut metagenome,” whereas this sample actually corresponded to a fecal transplant from a human to a germ-free mouse (ERR11510989). Another case involved an “air metagenome” from a swine facility that was incorrectly relabeled to “pig gut metagenome” (SRR13618035).

## 3 Discussion

In this work, we present three complementary resources for large-scale metagenomic comparison: (i) FracMin-Hash (FMH) sketch database, (ii) a hypervector database, and (iii) an all-vs-all similarity matrix, together with several applications that demonstrate their utility at different scales. The three resources serve distinct but complementary roles. FMH sketches are advantageous because they preserve sampled 31-mer identities, enabling sequence-aware analyses such as containment searches, taxonomic classification, average nucleotide identity estimation, and diversity analysis. However, performing large-scale similarity searches directly across millions of FMH sketches is computationally demanding. Hypervectors address this scalability challenge by sacrificing individual *k*-mer identities in favor of a considerably more compact representation. Though hypervectors can yield little biological meaning of their own, they enable rapid Jaccard similarity estimation, nearest-neighbor search, and large-scale sample comparison. The all-vs-all similarity matrix provides a complementary global view by precomputing relationships among the metagenomic experiments. Together, these representations allow users to move between sequence-level analysis, rapid similarity search, and archive-wide exploration depending on the question of interest.

The central contribution of this work, however, is not simply the sketching of millions of metagenomic datasets, but the transformation of the SRA metagenome collection into a resource that can be searched, compared, and explored directly at scale. By providing representations with different trade-offs among retained sequence information, storage requirements, and computational efficiency, together with a global similarity matrix, we make experiment-to-experiment comparison across millions of metagenomes computationally practical. In demonstrating the use cases of these resources, we place particular emphasis on applications enabled by large-scale experiment-to-experiment comparison, rather than on analyses that are already well established for sketch-based representations [27, 14, 15, 16, 31, 32].

As with any data reduction approach, it is important to quantify the error it introduces to the downstream estimates by characterizing their bias and/or giving confidence intervals [4]. For hypergen sketching [38], this has hitherto not been possible to do comprehensively due to the vast range of possible inputs. Here, we give a provably correct algorithm for sampling from the error distribution of Jaccard estimates from hypergen sketches, allowing us to show that the estimator had no bias and to compute confidence intervals for any input. We believe that this makes the hypergen sketch Jaccard estimator an attractive option for other applications besides the one in this paper.

Our large-scale view of the data also makes sequence similarity particularly useful as an independent signal for assessing the organization and quality of public metagenomic data. In particular, the identification of duplicated accessions and potential metadata inconsistencies illustrates how sequence-derived relationships can provide an additional layer of information beyond existing annotations. However, such findings should not be treated as definitive corrections, since apparent discrepancies may reflect legitimate biological or experimental differences. Any proposed re-annotation should therefore be supported by careful examination of the corresponding studies and experimental context. Nevertheless, the speed of hypervector comparison suggests a broader future role in systematic metadata harmonization. Such approaches may become particularly valuable as public sequencing archives are increasingly used for large-scale statistical and machine-learning applications, where metadata quality and consistency are critical.

Building on these possibilities, we anticipate that the public release of the FMH sketches, hypervectors, similarity matrix, and associated search tools will support applications beyond those demonstrated here, including environment-specific comparative studies, contamination and outlier detection, clustering-based biodiversity analyses, and recruitment of related datasets for downstream analysis. More broadly, we expect that making the metagenomic SRA directly navigable by sequence similarity will enable questions and workflows that are difficult to anticipate in advance.

## 4 Online methods

In Section 4.1, we start by defining key concepts and our Jaccard estimator. In Section 4.2, we prove the main theorem characterizing its distribution. In Section 4.3, we present an algorithm for sampling from this distribution and show how it can be used to measure the estimator”s bias and root mean square error, as well as how to construct prediction and confidence intervals. In Section 4.4, we describe the computational pipeline used to create the sketches. In Section 4.6 we detail the metadata relabeling algorithm. In Section 4.7, we describe the compression schemes we used.

### 4.1 Definitions

Our FracMinHash and hypergen sketches are controlled by two parameters, *p* ∈ [0, 1] and *d* ∈Z^+^, and defined in terms of two other hash functions, *f* and *g*. The function *f* maps a *k*-mer to a real number from the unit interval. The function *g* maps a *k*-mer to a *d*-dimension vector in {−1, +1}^*d*^. The FracMinHash sketch is defined as a function on a set of *k*-mers as FMH(*A*) = {*f* (*τ*) | *τ* ∈ *A, f* (*τ*) ≤ *p*}. Finally, *f* an d *g* are combined into the main hypergen sketch function *h* which is defined on a set of *k*-mers as *h*(*A*) = Σ*x*∈FMH(*A*) *g*(*x*). The function *h* is also referred to as the DotHash of the set FMH(*A*) or the hypergen sketch of *A*.

In the following, we will assume that *f* and *g* have the strongest possible randomness, which in this case means that they are 4^*k*^*-uniform*. A function is 4^*k*^*-uniform* if and only if 1) for any *k*-mer *x* and value *y* in the co-domain, the probability that *x* hashes to *y* is 4^−*k*^, and 2) for any *n* ≤ 4^*k*^ and any set of *n* distinct *k*-mers, their hash values are mutually independent. Note however that our practical implementations of *f* and *g* (described in Sup. Note 8) do not provably achieve this level of randomness, which may introduce an error which we evaluate separately in Sup. Note 8.1.

Given two sets *S*_1_ and *S*_2_, the Jaccard between them is 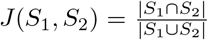. Given a vector *v*, we refer to the scalar in its *i*^th^ dimension as *v*_*i*_. Given a vector *v* and a scalar *x, v* − *x* is the vector obtained by subtracting *x* from every dimension of *v*. The *L*_2_ norm of *v* is denoted by ∥*v* _2_∥ and *u* · *v* refers to the dot product of vectors *u* and *v*.

For the sake of completeness, we recapitulate the derivation of the Jaccard estimator [25, 38]. Given two *k*-mer sets *A* and *B* and their respective hypervectors *h*(*A*) and *h*(*B*), the estimator is derived based on two approximations. The first is based on FracMinHash theory, which says that *J*(FMH(*A*), FMH(*B*)) is a nearly unbiased estimator of *J*(*A, B*) [27]. The second is based on DotHash theory [25], which states that for any two sets *S*_1_ and *S*_2_, *h*(*S*_1_) · *h*(*S*_2_) is an unbiased estimator of *d* · |*S*_1_ ∩ *S*_2_|. We then get

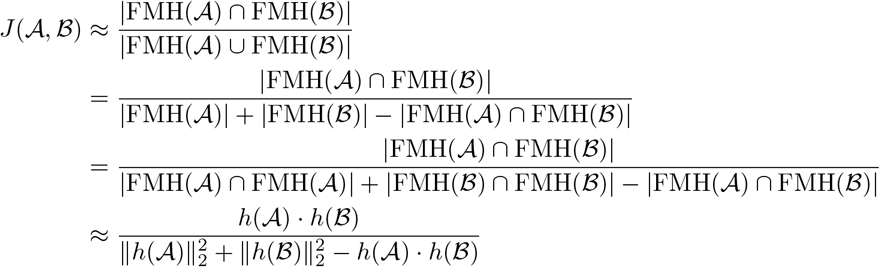

As a result, we define our estimator as

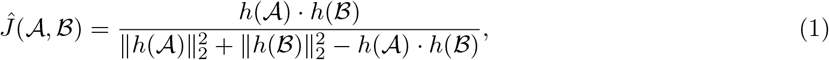

### 4.2 Main theorem

We now give a characterization of the distribution of *Ĵ*. First, we characterize the distribution on *h*(·) in terms of draws from Binomial distributions.

#### Lemma 1.

*Let C be any set of k-mers and let C* = *h*(*C*). *Let X* = |*FMH*(*C*)| *and let* 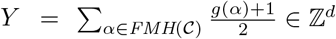. *Then*

1. *C* = 2*Y* − *X*.
2. *X* ~ Binom(|*C*|, *p*),
3. *conditioned on X* = *x, Y*_1_, … *Y*_*d*_ *is independently and identically distributed with Y*_*i*_ ~ Binom(*x*, 0.5).

*Proof*. To prove the first statement, 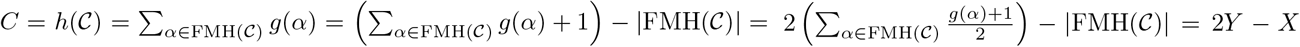. The second statement follows by definition of FMH and by the 4^*k*^-uniformity of *f*, which imply that every *k*-mer in *C* is included in FMH(*C*) with probability *p*, independent of other *k*-mers in *C*.

To prove the third statement, let *i* ∈ *{*1, … *d*}, and consider 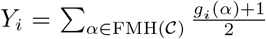. By definition of 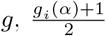 follows the Bernoulli distribution with success probability 0.5 and *g*_1_(*α*), …, *g*_*d*_(*α*) are mutually independent. Each *Y*_*i*_ is the sum of *X* mutually independent Bernoulli variables (with success probability 0.5). Conditioned on *X* = *x, Y*_*i*_ ~ Binom(*x*, 0.5) and *Y*_1_, …, *Y*_*d*_ are mutually independent. □

A key consequence of Lemma 1 is that the distribution of *h*(*C*) depends only on |*C*| and, once |*C*| is fixed, does not depend on the *k*-mers of *C*. This is crucial for simulations via sampling, as the results of a single simulation on a given size can apply to all *k*-mer sets having that size.

Next, we characterize the distribution of *Ĵ* (*A, B*), given two *k*-mers sets *A* and *B*. The difficulty is that the distributions of *h*(*A*) and *h*(*B*) are correlated if *A* and *B* share *k*-mers. Therefore, we cannot naively apply Lemma 1. However, if we partition *A* ∪ *B* into three non-overlapping sets *C*^1^ = *A \B, C*^2^ = *A* ∩ *B*, and *C*^3^ = *B \ A*, then the distributions *h*(*C*^1^), *h*(*C*^2^), and *h*(*C*^3^) are mutually independent. The distribution of *Ĵ*(*A, B*) can then reconstruction from these three distributions.

#### Theorem 1.

*Let A and B be two k-mer sets and let C*^1^ = *A \ B, C*^2^ = *A* ∩ *B,and C*^3^ = *B \ A. For all j* ∈ *{*1, 2, 3}, *let C*^*j*^ = *h*(*C*^*j*^), *let X*^*j*^ = |*FMH*(*C*^*j*^)| *and let* 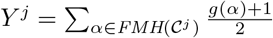. *Then*

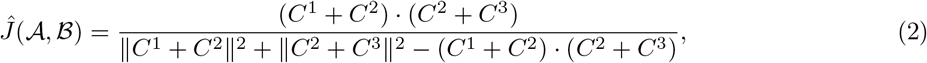

*and, for all j* ∈ *{*1, 2, 3}, *C*^*j*^ = 2*Y* ^*j*^ − *X*^*j*^, *where*

1. *X*^*j*^ ~ Binom(|*C*^*j*^ |, *p*),
2. 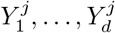*conditioned on X*^*j*^ = *x are i*.*i*.*d. with distribution* Binom(*x*, 0.5).
3. *for all* 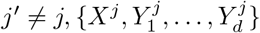*is independent of* 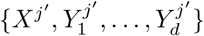.

*Proof*. By construction, on any disjoint sets *S*^1^ and *S*^2^, FMH(*S*^1^ ∪ *S*^2^) = FMH(*S*^1^)∪FMH(*S*^2^) and FMH(*S*^1^) and FMH(*S*^2^) are disjoint. As a consequence,

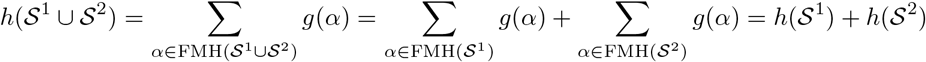

In particular, we can write *h*(*A*) = *C*^1^ + *C*^2^ and *h*(*B*) = *C*^2^ + *C*^3^. Together with the definition of *Ĵ* (Eq. (1)), this implies Eq. (2).

The fact that *C*^*j*^ = 2*Y* ^*j*^ −*X*^*j*^ and statements 1 and 2 follow directly from Lemma 1. To see statement 3, observe that given two distinct *k*-mers *τ* and *τ*^*′*^, *f* (*τ*) and *f* (*τ*^*′*^) are independent, and *g*(*τ*) and *g*(*τ*^*′*^) are independent. Therefore, for any two disjoint sets of *S*^1^ and *S*^2^, the random variables defined on them using *f* and *g* are also mutually independent.

A key consequence of Theorem 1 is that the distribution of *Ĵ* (*A, B*) only depends on |*A \ B*|, |*A* ∩ *B*|, |*B \ A*|and and, once these quantities are fixed, does not depend on the sets *A* and *B* themselves. This greatly reduces the space needed to be explored during simulations.

### 4.3 Error sampling algorithm and prediction and confidence intervals

We now present an algorithm for sampling from the error distribution of *Ĵ* and show its uses. Theorem 1 shows that for any *A* and *B*, the error depends on the size of the intersection and the sizes of the set differences. We therefore let *s*^1^ = |*A \ B*|, *s*^2^ = |*A* ∩ *B*|, and *s*^3^ = |*B \ A*| be the parameters to our algorithm.

The error of *Ĵ* is defined as Err(*A, B*) = *Ĵ* (*A, B*) − *J*(*A, B*). Since *J*(*A, B*) = *s*^2^*/*(*s*^1^ + *s*^2^ + *s*^3^), the distribution of Err(*A, B*) is simply the distribution of *Ĵ* (*A, B*) shifted by *s*^2^*/*(*s*^1^ + *s*^2^ + *s*^3^). Therefore, sampling from Err(*A, B*) is algorithmically equivalent to sampling from *Ĵ* (*A, B*).

Theorem 1 expresses *Ĵ* (*A, B*) as a function of random variables 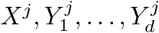, for *j* ∈ {1, 2, 3}. More importantly, it describes the dependence structure between these variables as well as their distributions. Namely, *X*^1^, *X*^2^, and *X*^3^ are mutually independent and Binomially distributed, and, after conditioning on the values of these three variables, all the *Y* variables become mutually independent and Binomially distributed. This immediately leads to an algorithm to sample from the distribution of Err(*A, B*). In particular, for each given *s*^1^, *s*^2^, and *s*^3^,

1. For each *j*, draw *x*^*j*^ ~ Binom(*s*^*j*^, *p*).
2. For each *j* and *i* ∈ *{*1, …, *d*}, draw *y*^*j*^ ~ Binom(*x*^*j*^, 0.5) and define *c*^*j*^ = 2*y*^*j*^ − *x*^*j*^.
3. Return 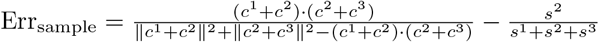

By repeatedly sampling using this algorithm, we can estimate the bias and root mean square error of our estimator. Crucially, since Theorem 1 states that the distribution of *Ĵ* (*A, B*) depends only on *s*^1^, *s*^2^, and *s*^3^, the space which the sampling needs to explore does not involve *k*-mer sets but only three integer values.

Our *Ĵ* estimator is a type of *point estimate*, meaning that it gives a single estimated value but does not capture its certainty. Prediction and confidence intervals allow us to go beyond that and capture certainty. Given parameters *α* ∈ (0, 1), *j* ∈ (0, 1), and non-negative integers *n*_*a*_ and *n*_*b*_, a *symmetric α-prediction interval* is the interval (*j* − *ϵ, j* + *ϵ*) such that for all *k*-mer sets *A* and *B* with

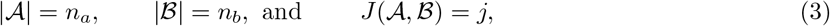

we have Pr[|*Ĵ* (*A, B*) − *j*| *< ϵ*] = *α*. Note that the three conditions in Eq. (3) are algebraically equivalent to

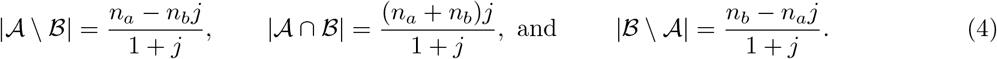

To compute the prediction interval, we run our algorithm to generate many samples using parameters 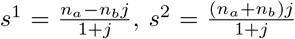, and 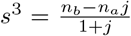. We then define *ϵ* as the smallest value such that at least an *α* fraction of the samples satisfy |Err_sample_| *< ϵ*.

Given two *k*-mer sets *A* and *B* and a parameter *α* ∈ (0, 1), an *α-confidence interval* is an interval (*j*_low_, *j*_high_) which contains *J*(*A, B*) with probability *α*. In our setting, we want to compute *j*_low_ and *j*_high_ only from |*A*|, |*B*|, and *Ĵ* (*A, B*). We can do so directly from the prediction intervals. In particular, we say that a value *j* qualifies for the confidence interval if the symmetric *α*-prediction interval around *j* contains *Ĵ* (*A, B*). We then let *j*_low_ and *j*_high_ be the interval that captures all candidate values *j*.

### 4.4 Computational pipeline

We now describe the computational pipeline used to generate our resources. The collection of *n* = 4,774,811 metagenomic accessions was identified by querying SRA metadata for accessions where either: (i) the `organism` field contained the substring “metagenome”, or (ii) the `librarysource` field was annotated as “METAGENOMIC” or “METATRANSCRIPTOMIC”. FracMinHash generation was performed on Amazon Web Services using AWS Batch and Docker containers executed on EC2 c5d-family Spot instances. Logan v1.1 unitig files were retrieved from Amazon S3 and decompressed onto the instance local storage. For each accession, Sourmash [16] “scripts manysketch” was run with scaled=1000, noabund, and k=15,31,33; only the k=31 sketches were used in the present study. Source code for running the cloud computation system is available at https://gitlab.pasteur.fr/rchikhi_pasteur/logan-analysis. To obtain the hypervector from an FMH sketch, we first mapped each FMH hash to a random 2048-dimensional hypervector with values of *{*+1, −1} and then summed the results (see Sup. Note 8 for details). These steps were all done in *O*(*n*) time.

To compute all-pairs estimates using Eq. (1), we need to compute the *L*_2_ norm of every hypervector (which takes *O* (*n*) total time) and the the dot product between all pairs of hypervectors (which takes *O* (*n*^2^) total time). We optimize the second step using two approaches. The first is that the all-versus-all dot product computation is done using the powerful Eigen `C++` linear algebra library [13]. The second is that we partition the sorted list of accession IDs into equally-sized shards, and, for each shard, compute the dot products between the shard”s accessions and the complete set of accessions. In other words, we divide the computation of the matrix into consecutive blocks of rows and compute each block separately. This reduces memory and allows parallel computation of the matrix.

### 4.5 Co-assembly

The eight targets with project helpers were selected from distinct BioProjects with low estimated coverage (median Logan contig coverage ≤4x among the first 1,000 contigs) for which the similarity of the SRA-wide helper exceeded that of the project helper by at least 0.02. Targets were ranked by decreasing similarity advantage.

Eligible helpers were required to have paired-end Illumina WGS metagenomic reads, at least 200 million sequenced bases. For each target, helpers were selected as the highest estimated-Jaccard eligible helper within the target BioProject and outside of it, using the top 1,000 nearest-neighbor search results. Assemblies were performed using MEGAHIT v1.2.9 with the `meta-sensitive` preset [20].

### 4.6 Metadata correction

For our proof-of-concept metadata relabeling algorithm, we first need to account for the redundancy in many of the possible `organism` labels. We therefore manually partition them, with each partition group corresponding to likely similar communities, e.g., “human feces” and “human gut metagenome” (Sup. Table 2). To decide if an accession *A* should be relabeled, we select one accession from every BioProject in *A*”s similarity neighborhood, choosing the one that is a neighbor of *A* with the highest similarity to *A*. The selected accessions are further filtered to remove those with less than 10% similarity to *A* and those with the generic tags of “metagenome”, “human metagenome”, and “Homo sapiens.” If at least four and at least 80% of tags of the selected accessions belong to a partition group different from that of *A*”s tag, we propose that group”s representative as a relabeling candidate for *A*.

### 4.7 Compression

In order to reduce the space of our sketch databases, we developed custom compression algorithms. To compress the FMH sketch database, we rely on Elias-Fano encoding [9, 11]. Elias-Fano exploits the fact that the total number of distinct 64-bit hashes observed in our database (*N* = 33 × 10^9^) is much less than the space of possible 64-bit integers (~2 × 10^19^) and, moreover, the hashes within a sketch can be stored in sorted order without any loss of information. To implement the compression, we first create a sorted global array of the *N* hashes. We then relabel each 64-bit hash in the database with an identifier in {0, …, *N*− 1}, corresponding to its rank in the global array. We then apply Elias-Fano to individually compress both the global array and each accession”s relabeled sketch. Finally, we run the generic `xz` archival compression on these files. Sup. Note 7 contains further details.

To compress our similarity matrix, we exploit its sparsity. First, we create a hash table *H* assigning each accession ID to its rank in the sorted list of accession IDs, i.e. to its row index in the similarity matrix. We encode the non-zero values of a row *r* using two vectors, *N*_*r*_ and *J*_*r*_. The *N*_*r*_ list stores an ascending list of the indices of the non-zero columns, using Rice encoding [23]. The *J*_*r*_ list stores the corresponding Jaccard estimate values, with each value quantized into 256 bins and stored in a byte.

Recall that our matrix”s rows are partitioned into shards, corresponding to consecutive blocks of rows. For each shard, we store the *N*_*r*_ and *J*_*r*_ lists jointly interleaved in a file. In order to allow random access to the *i*^th^ row within the shard, we store an additional offset array *O* for each shard. In particular, *O*[*i*] is the offset in the joint list corresponding to the start of *N*_*r*_, where *r* is the *i*^th^ row in the shard. *O* is encoded using delta encoding [23], which exploits that its values are increasing. This compression scheme allows us to map a query accession to its row index *r* and its corresponding shard and then use *O* to locate *N*_*r*_ and *J*_*r*_ which can then be decompressed and reported to the user.

## Acknowledgments

This work was was supported by NIGMS/NIH grant R01GM146462. R.F. was supported by the European Union Horizon 2020 research and innovation program No. 101088572 (ERC IndexThePlanet). This work was initiated during a visit of R.F. to Penn State which was supported by a grant from the European Union Horizon 2020 research and innovation program No. 872539 (PANGAIA). The authors acknowledge the Texas Advanced Computing Center (TACC) at The University of Texas at Austin for providing computational resources that have contributed to the research results reported within this paper.

## Supplementary Notes

### Supplementary Note 1: Data summary statistics

**Supplementary Table 1:**
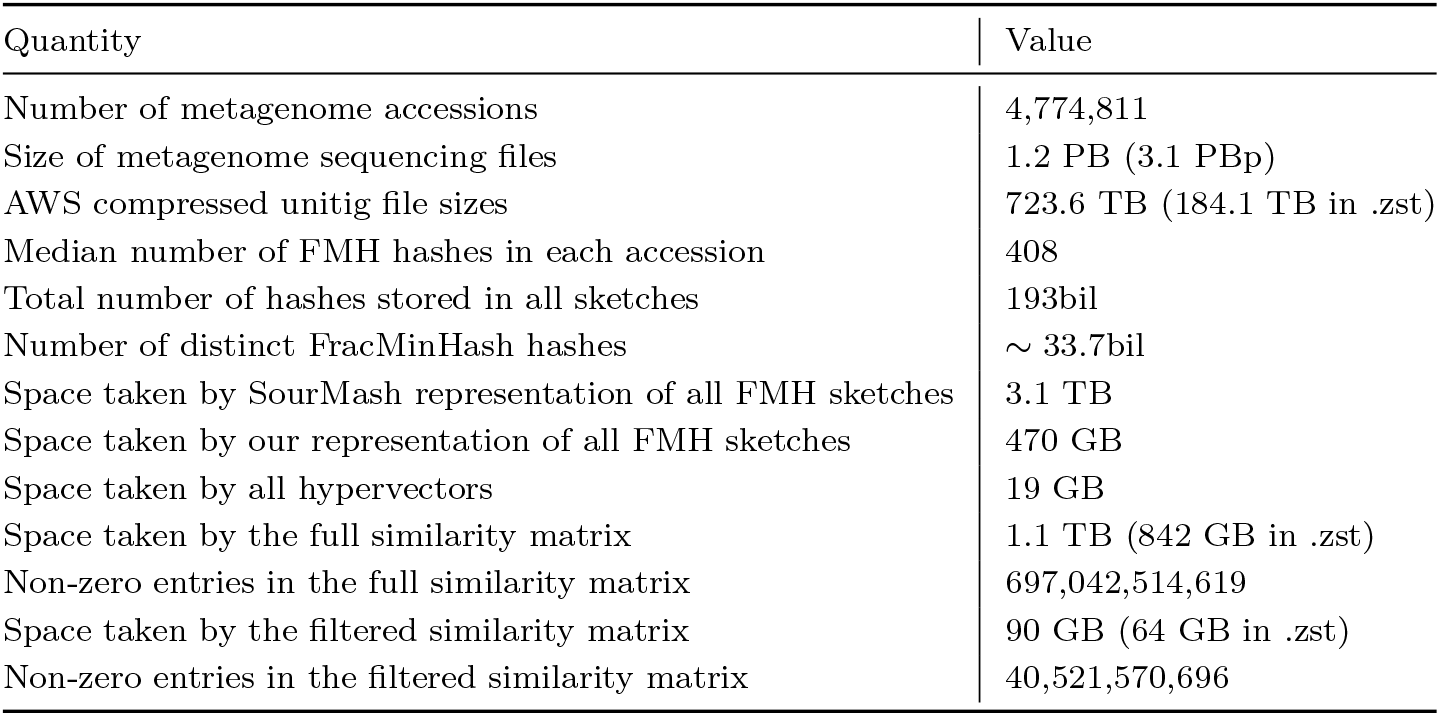
Data summary statistics.

| Quantity | Value |
| --- | --- |
| Number of metagenome accessions | 4,774,811 |
| Size of metagenome sequencing files | 1.2 PB (3.1 PBp) |
| AWS compressed unitig file sizes | 723.6 TB (184.1 TB in .zst) |
| Median number of FMH hashes in each accession | 408 |
| Total number of hashes stored in all sketches | 193bil |
| Number of distinct FracMinHash hashes | $\sim 33.7$bil |
| Space taken by SourMash representation of all FMH sketches | 3.1 TB |
| Space taken by our representation of all FMH sketches | 470 GB |
| Space taken by all hypervectors | 19 GB |
| Space taken by the full similarity matrix | 1.1 TB (842 GB in .zst) |
| Non-zero entries in the full similarity matrix | 697,042,514,619 |
| Space taken by the filtered similarity matrix | 90 GB (64 GB in .zst) |
| Non-zero entries in the filtered similarity matrix | 40,521,570,696 |

### Supplementary Note 2: Newly observed 31-mers for soil and human gut metagenomes

**Supplementary Figure 1:**
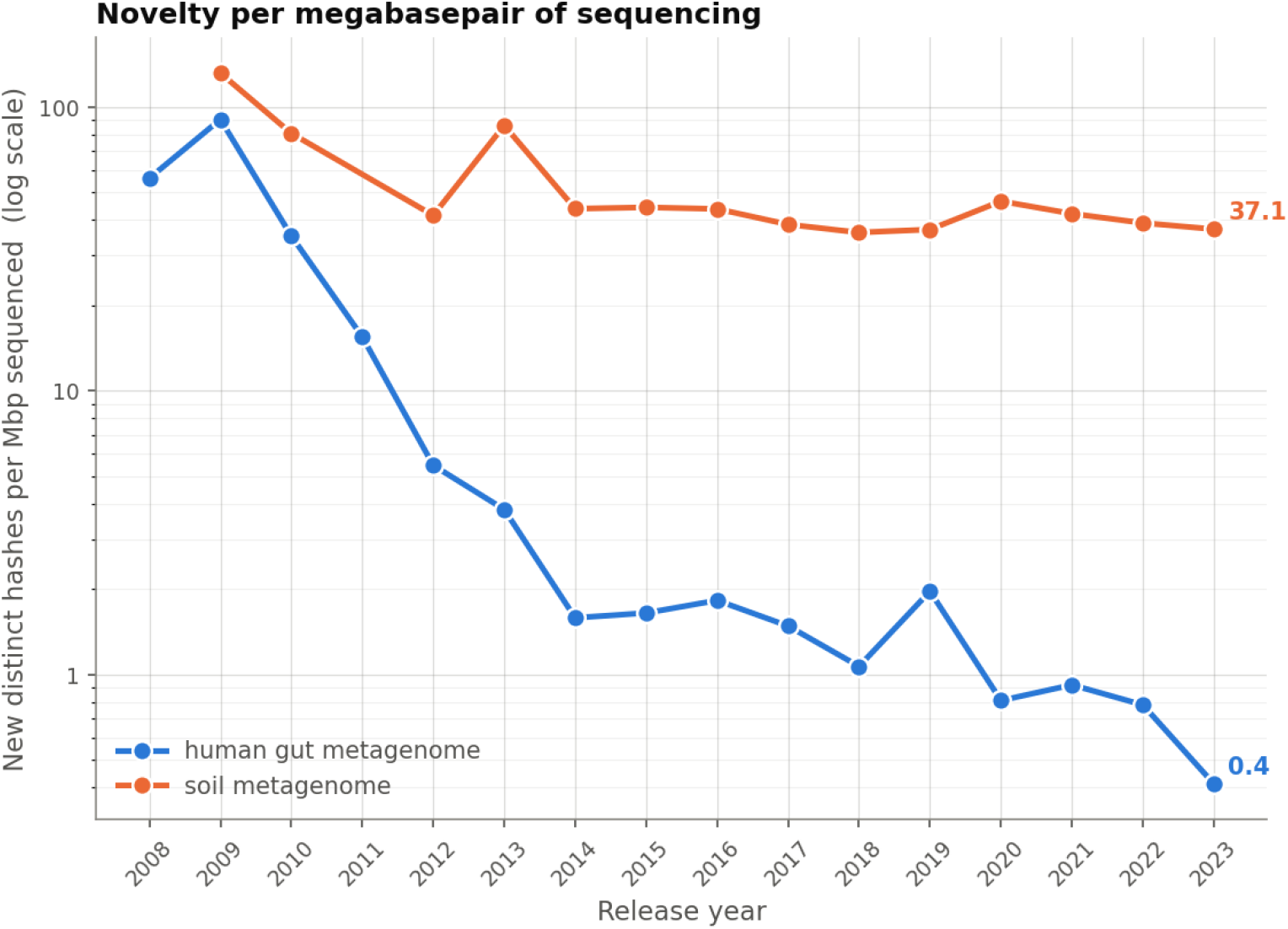
Novel 31-mer hashes returned per megabasepair of sequencing, 2008–2023, for soil and human gut WGS metagenomes. The log-scale *y*-axis is the number of such hashes appearing in that environment”s WGS runs released that year, divided by the total megabasepairs those runs contributed. Soil is essentially flat across the last decade, from 43.8 novel hashes per Mbp in 2014 to 37.1 in 2023, whereas the human gut rate falls from 1.58 to 0.41 over the same period. By 2023 a megabasepair of soil sequencing returned roughly 90 times more previously-unseen sequence than a megabasepair of human gut sequencing, despite the human gut receiving its largest and deepest sequencing year on record in 2023 (42,098 runs, 188.9 Tbp). Note that the plot does not require a *k*-mer to appear in multiple years (unlike Figure 2(c,d)), because the focus is on relative, rather than absolute, trajectories.

### Supplementary Note 3: Source of error in our Jaccard estimator

**Supplementary Figure 2:**
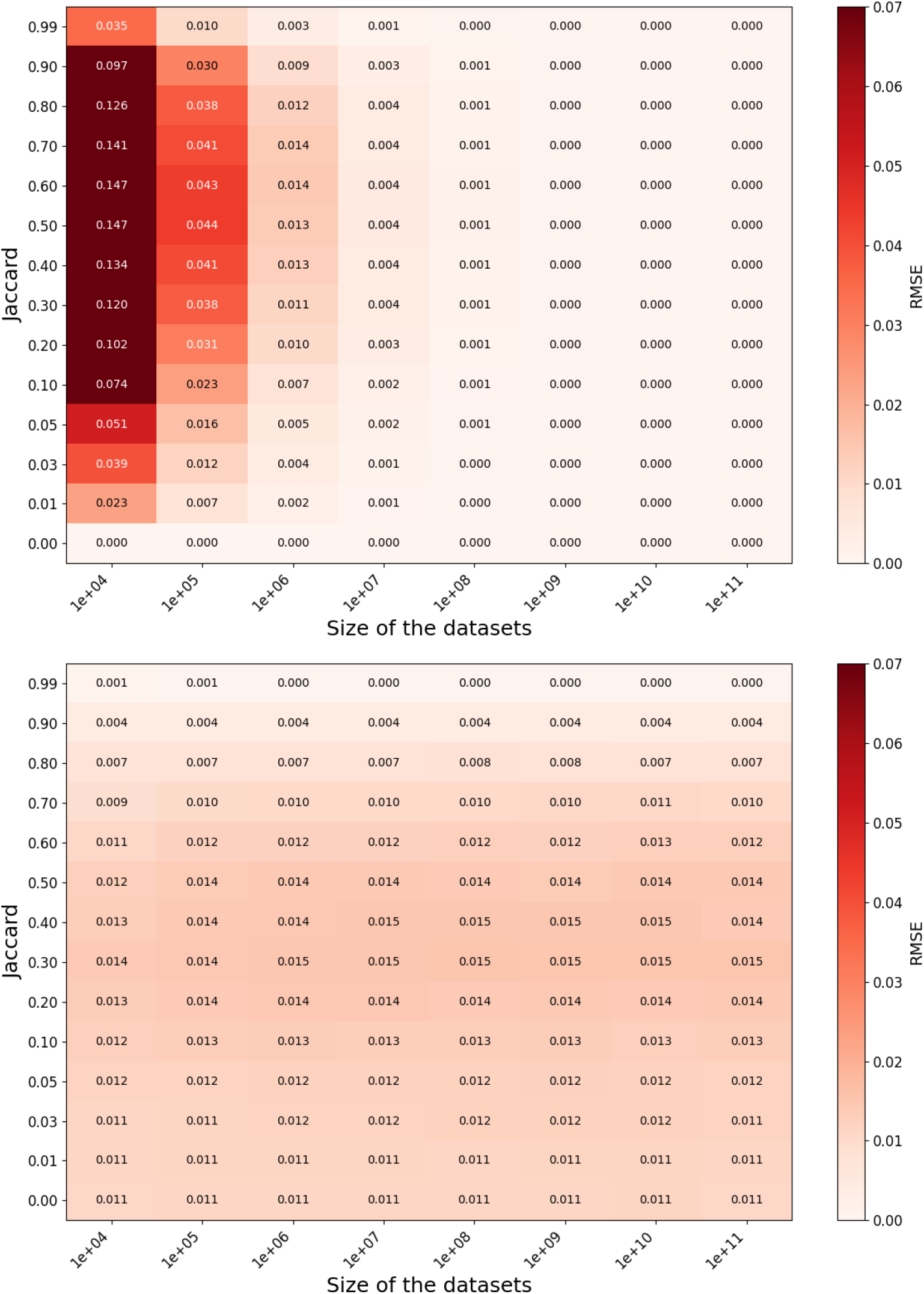
Decomposition of the error of our Jaccard estimator into FracMinHash (top) and Dothash (bottom) errors. The error is evaluated on pairs of datasets of equal sizes and is reported as the root mean square error (RMSE).

### Supplementary Note 4: Metadata correction

**Supplementary Table 2:** Groups of equivalent organism tags, as used in Section 2.3.2. All `organism` tags not mentioned in this table were assigned to a group containing only themselves.

| Group Representative | Additional Group Tags |
| --- | --- |
| general metagenome | human metagenome, metagenome, Homo sapiens |
| human blood metagenome | blood metagenome |
| human gut metagenome | gut metagenome, feces metagenome, human feces metagenome |
| human lung metagenome | lung metagenome, respiratory tract metagenome, upper respiratory tract metagenome, human tracheal metagenome |
| human oral metagenome | oral metagenome, human saliva metagenome |
| human skin metagenome | skin metagenome |
| human urinary tract metagenome | urine metagenome |
| human vaginal metagenome | vaginal metagenome |
| hypersaline metagenome | hypersaline lake metagenome, salt marsh metagenome, salt lake metagenome |
| sludge metagenome | activated sludge metagenome, bioreactor metagenome, bioreactor sludge metagenome, anaerobic digester metagenome |
| unidentified | unidentified, uncultured bacterium, uncultured virus, uncultured crAssphage |
| bovine gut metagenome | bovine metagenome |
| canine gut metagenome | canine metagenome |
| fish gut metagenome | fish metagenome |
| insect gut metagenome | insect metagenome |
| mouse gut metagenome | mouse metagenome, Mus musculus, Rattus norvegicus |
| pig gut metagenome | pig metagenome |
| sheep gut metagenome | sheep metagenome |
| dust metagenome | indoor metagenome |
| freshwater metagenome | lake water metagenome, riverine metagenome, drinking water metagenome, groundwater metagenome |
| hot springs metagenome | hydrothermal vent metagenome |
| marine metagenome | seawater metagenome, marine plankton metagenome |
| manure metagenome | poultry litter metagenome |
| sediment metagenome | freshwater sediment metagenome, marine sediment metagenome |
| soil metagenome | rhizosphere metagenome, root metagenome, terrestrial metagenome |
| viral metagenome | human viral metagenome, aquatic viral metagenome, phage metagenome |
| Equus caballus | Equus asinus asinus |
| Lassa virus | Mammarenavirus lassaense |
| Human rhinovirus | Human rhinovirus sp. |
| SARS-CoV-2 | Severe acute respiratory syndrome coronavirus 2 |
| Sinorhizobium | Sinorhizobium meliloti |
| Treponema pallidum | Treponema pallidum subsp. pallidum SS14, Treponema pallidum subsp. pallidum, Treponema pallidum subsp. pertenue |

**Supplementary Table 3:**
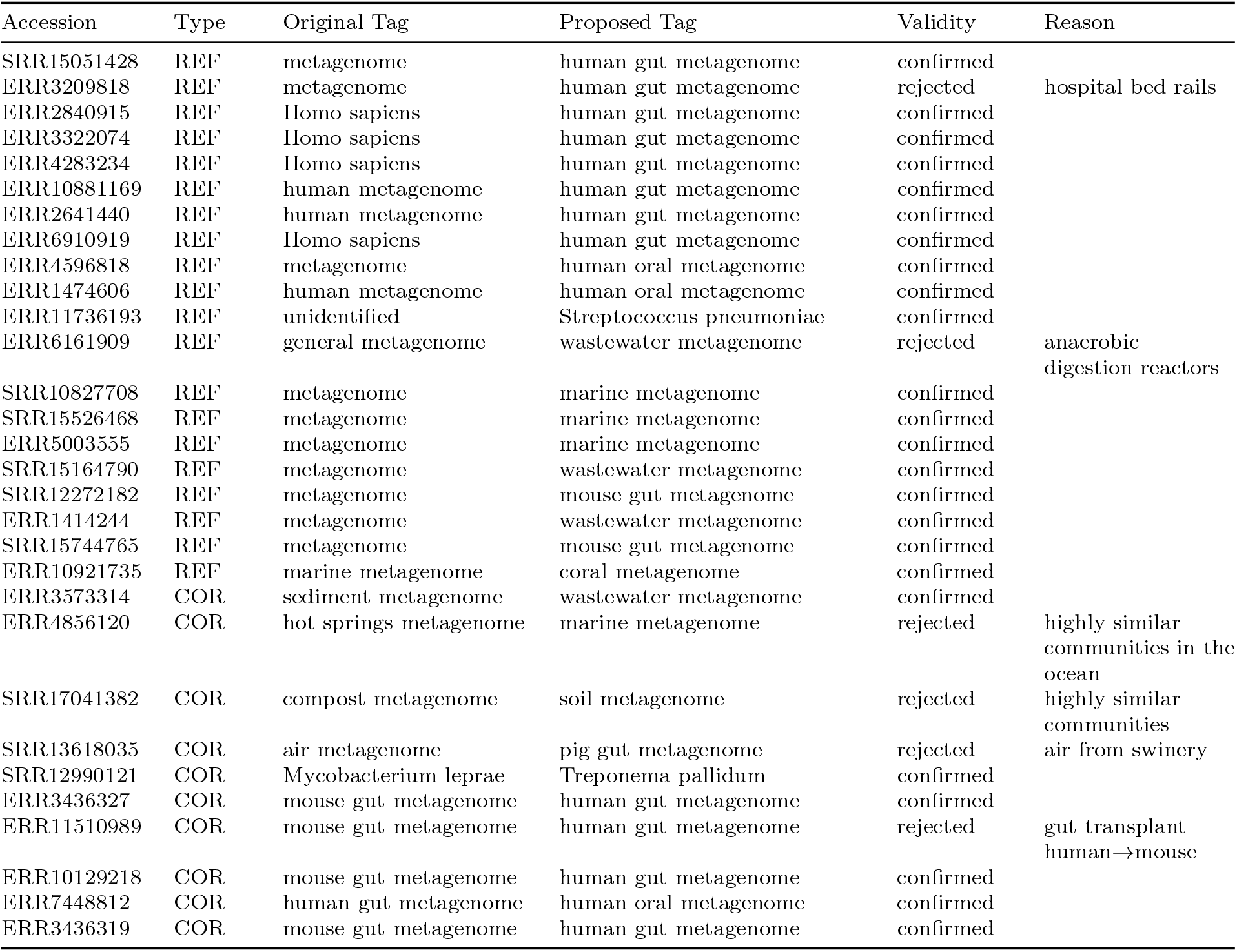
Spot-checks of 30 proposed organism tag relabelings. The “Type” column refers to the type of relabeling, which is either a refinement (REF) or correction (COR) of the existing label.

### Supplementary Note 5: Filtering the similarity matrix

**Supplementary Table 4:** Statistics of the similarity matrices under different dataset variations. The first column lists the matrix type: “Full” includes all samples, “WGS only” includes only whole-genome sequencing samples, and “Non-WGS only” includes only non-WGS samples. “≥ *N* FMH hashes only” denotes matrices constructed from samples containing at least *N* FracMinHash signatures. “*s* Jaccard threshold” correspond to matrices including only neighbors with ≥ *s* Jaccard similarity. The last row is the strategy we ultimately chose for our filtered matrix: for an accession X, a neighbor Y with similarity *s* is retained if *s* ≥ 0.2 or fewer than 10, 000 neighbors of X have similarity greater than *s*.

| Type | Size (GB) | N. samples | N. non-zero entries | Mean n. of neighbors | Median n. of neighbors |
| --- | --- | --- | --- | --- | --- |
| Full | 1,147 | 4,774,811 | 697,037,739,808 | 145,982 | 58,043 |
| WGS only | 26 | 600,700 | 17,361,325,130 | 28,902 | 2,479 |
| Non-WGS only | 1,071 | 4,174,111 | 661,685,289,600 | 158,521 | 74,329 |
| $\geq 1,000$ FMH hashes only | 92 | 1,252,002 | 57,480,996,916 | 45,911 | 10,446 |
| $\geq 500$ FMH hashes only | 314 | 2,097,581 | 198,800,016,376 | 94,776 | 33,668 |
| $\geq 100$ FMH hashes only | 988 | 3,976,546 | 612,903,068,344 | 154,130 | 70,144 |
| $\geq 10$ FMH hashes only | 1,141 | 4,676,975 | 695,078,819,348 | 148,617 | 62,634 |
| 0.2 Jaccard threshold | 10 | 4,774,811 | 4,000,916,936 | 838 | 75 |
| 0.1 Jaccard threshold | 227 | 4,774,811 | 120,142,393,410 | 25,162 | 3,809 |
| Top 10,000 neighbors / Jaccard $\geq 0.2$ | 90 | 4,774,811 | 40,516,795,885 | 8,486 | 10,518 |

### Supplementary Note 6: Experimental details

#### 6.1 k-mer diversity and novelty

Figure 1 shows how much previously-unseen sequence an environment still returns per unit of sequencing, for the soil and human gut metagenomes. To measure this, we partitioned the corpus by the SRA `organism` metadata field, taken verbatim without harmonization of near-synonyms, and restricted attention to runs with `assay type=WGS`, and a non-null release date. This yields 165,335 runs (577.7 Tbp) for `human gut metagenome` and 20,723 runs (174.7 Tbp) for `soil metagenome`. A hash is counted as *new* in year *Y* if *Y* is the first calendar year in which it appears anywhere in the corpus. We counted these new hashes per environment and divided by the total megabasepairs sequenced across those runs (the SRA `mbases` field, complete for both strata). Note that the resulting rate is not normalised for the fact that the two environments were sequenced at very different scales; this is deliberate, since progressive exhaustion of an environment is precisely a consequence of how heavily it has already been sampled.

#### 6.2 Error analysis

##### Bias

We evaluated 198 distinct Venn configurations, spanning *k*-mer set sizes from 10^4^ to 10^10^ and Jaccard similarities ranging from 0 to 0.99. For each configuration, we performed 30,000 independent estimations and measured the difference between the true Jaccard and the average value of the estimator.

##### RMSE measurement

We selected 13 Jaccard values ranging from 0 to 0.99 and 14 dataset sizes ranging from 10^4^ to 10^10^. For each possible configuration of Jaccard and pairs of dataset sizes, we performed 30,000 independent estimations, with which used for downstream error analysis shown in Figure 3(c,d) and Sup. Fig. 2.

##### Similarity detection false positive rate

To determine the 99.999% prediction interval around Jaccard of zero, we ran 500,000 simulations for each configuration. To run such a large number of simulations, we restricted ourselves to pairs of datasets of the same size.

### 6.3 Runtime benchmark of Jaccard estimators

Figure 3(b) shows the all-versus-all running times for computing Jaccard similarities across the first 3,536 SRA accessions (sorted alphabetically), using four different tools: `BinDash` v2.5 [39], `sourmash` v2.4.9 [16], `Dashing` v2.1.20 [2], and the hypervector Jaccard estimator we implement. All runs were performed single-threaded on an Intel Core Ultra 9 CPU. For `sourmash`, we used the same parameters as in the rest of the paper (a sampling rate of 1*/*1000). For *Dashing2* and *BinDash*, we used the default parameters but we set the sketch sizes to a fixed size of 4,096 bytes, matching the hypervector dimensions. We emphasize that this comparison is not intended as a rigorous benchmark, as the methods are not directly comparable: `sourmash` relies on variable-sized FracMinHash sketches, whereas the other tools use fixed-size sketches, and their error distributions differ. Speed vs. accuracy tradeoffs are possible using all methods and we do not explore them in depth. This experiment simply illustrates the computational efficiency of our hypervector-based all-versus-all Jaccard similarity approach and was used as preliminary analysis to determine the choice of software to use.

### Supplementary Note 7: Compression of FracMinHash sketch database

The complete FracMinHash sketch collection consisted of 4,774,811 `sourmash` signature files stored in `.sig` format, with a total uncompressed size of 3.1 TB. To make this dataset more practical to store and transfer, we developed a compression pipeline. The compression procedure consists of the following steps.

1. Construction of a global hash table: We first collected all FracMinHash hash values from the input `.sig` files and computed their global union. The union was sorted and deduplicated to produce a single ordered array of unique 64-bit hash values. Let *N* denote the total number of unique hashes, which in our case was 33.9B. Each hash was assigned a dense identifier from 1 to *N* based on its position in this sorted array. Therefore, the sorted binary `uint64` array serves as the global hash table, allowing the original hash value to be recovered from its identifier without storing an explicit key-value mapping.
2. Extraction of per-sketch hashes and metadata: For each `sourmash` signature file, we extracted the FracMinHash values from the `mins` field and stored them separately as a binary `uint64` file. The remaining signature information, including sketch parameters and metadata, was saved in a separate metadata directory after clearing the `mins` field, preserving the information needed for later signature reconstruction.
3. Conversion of hash values to dense identifiers: Each per-sketch hash list was then converted to the corresponding list of dense identifiers using the sorted global hash table. Because the global hash table is sorted, this conversion can be performed efficiently by searching for each hash value in the global array. The resulting identifier list represents the same FracMinHash sketch, but uses values from 1 to *N* rather than the original 64-bit hash values.
4. Compression using Elias-Fano encoding: The per-sketch identifier lists and the global hash table were compressed using Elias-Fano encoding [36]. Elias-Fano is a compression technique that stores a set of *u* integers from a universe of size *U* using 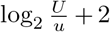 bits per integer. For an accession with *n* hashes, its per-sketch identifier list takes *n*(37 − log_2_ *n*) bits, given that *U* = 33.9B. The global hash table, where *u* = 33.9B and *U* = 1.84 × 10^16^, is predicted to take 89 GB. We implemented Elias–Fano encoding using the `bits` succinct data-structure library [12].
5. Final archive compression: Finally, the Elias-Fano compressed per-sketch identifier files, the Elias-Fano compressed global hash table, and the metadata directory were combined and compressed using `xz`.

After compression, the Elias-Fano encoded per-sketch hash identifier files occupied 390 GB, the compressed global hash table occupied 84 GB, and the separated metadata occupied 1.6 GB. The combined size of these components was 475 GB, which was further reduced to 470 GB after `xz` archival compression. Overall, the final compressed database was only 15% of the original `.sig` directory size, corresponding to an approximately 6.7*×* reduction in storage.

### Supplementary Note 8: Hash function implementation details

The FracMinHash (FMH) hashes were generated using the default hash function and seed of Sourmash [16], which is MurmurHash3 x64 128 low64 with a seed of 42. To compute hypergen sketches, recall that we need a function *g* to map from a 64-bit FMH hashcode to a bitvectors of size *d* = 2048. We implemented *g* using SplitMix [37, 34], a fast hash function that maps a 64-bit integer to a 64-bit integer that, in practice, mimics full randomness [6]. Let *x* be a 64-bit FMH hashcode. In our case, *g*(*x*) is 2048 bits, or, equivalently, 32 64-bit integers. To obtain *g*(*x*), we therefore return the concatenation of 32 SplitMix values, i.e. SplitMix(32*x*) · SplitMix(32*x* + 1) · SplitMix(32*x* + 31). Done in this manner, we guarantee that each any two distinct hashcodes will draw from different random bits. Moreover, our FMH sampling rate of 0.001 guarantees that *x* is always less than 2^64^·0.001 and hence 32*x* + 31 never exceeds 64 bits.

When computing a hypervector of a set of FMH hashcodes (i.e. *h*(*A*)), we sum the hypervectors of its constituent hashcodes, treating each 0 implicitly as a −1. To store the result, we use a 16-bit integer to store the value of each dimension. We find that 16 bits are sufficient, as the values stored grow with the square root of the number of FMH hashcodes. For example, a dataset containing a billion 31-mers would have a sketch of one million FMH hashcodes, and hence we expect the absolute values of the vectors to be on average 1000, well below the 2^16^ limit.

### 8.1 Accuracy of Theorem 1

Our confidence intervals rely on Theorem 1 to sample estimator error, which assumes that the underlying hash functions *f* and *g* are 4^*k*^-uniform. This is a theoretical ideal of randomness that is not achieved by hash functions in practice. However, since the hash implementations we use are known for having good random properties, we expect our error sampling to be nevertheless accurate. To validate this, we compare our estimator”s empirical error distribution with that obtained by the sampling algorithm of Section 4.3.

For each pair of accessions, the error is computed simply as the difference between the true Jaccard (using unitigs) and the estimated Jaccard (using Eq. (1)). However, we are limited in the number of pairs we could sample, as downloading unitigs and computing the Jaccard between them is time consuming. Moreover, we cannot draw pairs uniformly at random, as these are heavily enriched for accessions with Jaccard similarity near 0, whereas we wanted the samples to cover a broad range of *s*^1^, *s*^2^, and *s*^3^.

We therefore take a two stage approach. In the first stage, we create a smaller set of of pairs which is depleted of pairs with zero Jaccard similarity. To do so, we sample pairs uniformly until we can identify and retain 10,000 pairs with *Ĵ* ≥ 0.02. This depleted set creates a truncation bias, i.e. when the true Jaccard lies near 0.02, then only overestimates (e.g., *Ĵ* = 0.03) are retained, while underestimates (e.g., *Ĵ* = 0.01) are discarded. Therefore, in the second stage, we compute the ground-truth Jaccard on the depleted set and further exclude pairs with Jaccard less than 0.07. This removes the truncation bias, leaving 724 pairs with computed true Jaccard that we could use for full validation.

For each of the 724 (*A, B*) pairs, we start with an already-computed true and estimated Jaccard. Then, we make 10,000 draws from the distribution defined by Theorem 1 to obtain a distribution of Err(*A, B*). We then recorded the decile rank of the error of the estimated Jaccard within this simulated distribution. If the distribution given by Theorem 1 is correct, then these ranks should be uniformly distributed across the ten deciles. For our 724 pairs, the number of pairs in each of the ten deciles was [57, 68, 70, 79, 61, 73, 83, 85, 74, 74]. The chi-square test could show no deviation between this empirical distribution and a uniform one (p-value of 0.36). This test confirms that the empirical error of our estimator is consistent with the error computed using simulations in Section 4.3.

## Footnotes

1 https://github.com/RolandFaure/SRA-metagenome-sketches-databases

2 Because *k*-mer hash collisions are extremely rare, for the purposes of this analysis we equate the number of hashes with the number of canonical 31-mers

## Notes

https://github.com/RolandFaure/SRA-metagenome-sketches-databases

### Competing Interest Statement

The authors have declared no competing interest.

